# A translatable evaluation tool to study the biodistribution of clinically-available doxorubicin liposomes: PET imaging of [^89^Zr]Zr-Doxil and [^89^Zr]Zr-Talidox

**DOI:** 10.1101/2025.06.17.660144

**Authors:** Amaia Carrascal-Miniño, Aishwarya Mishra, Peter Gawne, Shane Angoh, Juliette Chupin, Jana Kim, Vittorio de Santis, Truc Pham, Fred Bark, Azalea Khan, Stefan Halbherr, Nicholas J. Long, Rafael T. M. de Rosales

## Abstract

**INTRODUCTION:** Doxil/Caelyx is a PEGylated liposomal formulation of the chemotherapeutic doxorubicin used in the clinic for Kaposi’s sarcoma, advanced ovarian cancer, progressive multiple myeloma and metastatic breast cancer. Talidox^®^, a smaller doxorubicin PEGylated liposome is undergoing clinical trials and has been proposed as an improvement on previous liposomal formulations for the treatment of advanced solid tumors. We aimed to validate an easily translatable radiolabeling method using zirconium-89 (^89^Zr) that enables quantitative whole-body PET imaging of these formulations to study their biodistribution and pharmacokinetics.

**METHODS:** [^89^Zr][Zr(oxinate)_4_] was produced using a kit-based approach followed by use as a direct radiolabeling agent of the liposomal formulations. DFT studies were performed to elucidate the mechanism behind the radiolabeling stability observed within the liposomes. Purified ^89^Zr-labelled Doxil/Talidox® liposomes (5 mg/kg doxorubicin dose) were administered in female BALB/c mice bearing 4T1 tumors. PET/CT imaging was acquired at 20 min, 24 h, 48 h, and 72 h, followed by post-mortem biodistribution at 72 h.

**RESULTS and DISCUSSION:** Both formulations were radiolabeled efficiently with high stability in serum *in vitro* for 72 h. *In vivo*, both formulations showed high tumor uptake at 72 h (18.5 ± 2.4 % IA/g for Doxil and 20.2 ± 2.3 % IA/g for Talidox). In general, *ex vivo* biodistribution showed similar uptake values for both formulations with high spleen/liver uptake and low bone uptake, confirming stability. Talidox® showed significantly lower spleen uptake and higher uptake in bone than Doxil. DFT studies confirmed that doxorubicin can form complexes with ^89^Zr that are more stable than [^89^Zr][Zr(oxinate)_4_], explaining the radiolabeling mechanism and stability results *in vitro* and *in vivo*.

**CONCLUSIONS:** Clinically available PEGylated liposomes containing doxorubicin can be efficiently radiolabelled with ^89^Zr for PET imaging studies, using a clinically translatable radiolabelling method.

**Highlights:**

- Doxorubicin-containing liposomes can be labeled with the positron-emitting radionuclide ^89^Zr with no impact on their original physicochemical properties.
- Radiolabeling is stable *in vivo* and enables imaging and biodistribution studies of the liposomes using positron emission tomography (PET).
- The radiolabeling method is clinically translatable and would allow early assessment of existing and novel doxorubicin liposome biodistribution in humans or personalized medicine (nanotheranostic) approaches.

## 1. Introduction

Doxorubicin, an anthracycline chemotherapeutic agent, has been employed in the management of various malignancies, including acute leukemia, Hodgkin’s and non-Hodgkin’s lymphoma, soft tissue sarcoma, and solid tumors such as breast cancer [1]. Despite the demonstrated efficacy, the utility of doxorubicin is limited by associated common adverse effects, notably cardiotoxicity, and bone marrow suppression, significantly compromising the patient’s quality of life.

Encapsulation of doxorubicin within liposomal delivery systems can enhance its tolerability and minimize toxicity. The first liposomal doxorubicin formulation was introduced to the market as Doxil^®^ in 1995 [2], this PEGylated liposome formulation is approved for the treatment of AIDS-related Kaposi’s sarcoma, ovarian cancer, multiple myeloma, and breast cancer [1]. Although with fewer reported side effects, the formulation does produce specific side effects linked with dosage, such as the hand-foot syndrome, a dose limiting side effect of the treatment [1] [3].

Talidox^®^ represents a smaller PEGylated liposomal formulation also containing doxorubicin which underwent a Phase I/IIa clinical trial in Switzerland (NCT03387917) for advanced solid tumors. This reduced size is designed to enhance its tumor permeation capacity [4] [5]. In the clinical studies, Talidox^®^ showed improved efficacy in advanced solid tumors and reduced side effects in comparison with other doxorubicin PEGylated liposomal formulations[5].

The mechanism underlying both PEGylated liposome formulations centers on facilitating doxorubicin delivery to the tumor via the Enhanced Permeability and Retention (EPR) effect. In brief, liposomes preferentially accumulate in well-vascularized tumors; the mechanism appears to be based on highly permeable capillaries that allow the extravasation of nano-scale agents into the tumor microenvironment. The latest research shows that the process is complex and active [6, 7]. Approximately a fifth of the endothelial cells appear to transport the nanoparticles into the tumor microenvironment, and these cells are upregulated in genes that facilitate nanoparticle transfer [6] [7]. This process is further enhanced by a reduced lymphatic clearance, leading to the accumulation of liposomal doxorubicin. The liposome must possess a relatively prolonged circulation half-life to achieve high accumulation, which is attained by PEGylation of the membrane [8].

Nonetheless, the EPR effect has been established as heterogeneous in humans [9]. Tumors exhibiting inadequate accumulation tend to exhibit suboptimal responses to Doxil and other liposomal formulations [10]. Ascertaining poor accumulation by monitoring the tumor’s response could potentially delay transitioning to more conventional therapeutic modalities. A non-invasive and quantifiable technique able to assess the accumulation of liposomes and predict the therapy outcome before a response is measured could improve the patient’s outcome, assess the effect of coadjuvant therapies, or detect resistance.

Nuclear imaging represents an excellent technique for this purpose due to its quantifiable and non-invasive nature. An array of SPECT (^111^In, ^99m^Tc) and PET radionuclides (^124^I, ^64^Cu) [10] have been used clinically as reviewed by Man *et al.* [10]. The radionuclide of choice should have certain characteristics for optimal clinical use. Firstly, PET imaging using positron-emitting radionuclides (^18^F, ^68^Ga, ^89^Zr) is superior to SPECT imaging (^111^In, ^99m^Tc) in quantification and sensitivity. Secondly, the chemical nature of the radionuclide will also affect radiolabeling stability and its half-life. Because of similar biodistribution and kinetics to antibodies (Abs), liposomes require radiolabeling with a radionuclide with a medium half-life, allowing imaging for at least 3 days. Good manufacturing practices (GMP) grade radionuclide availability must also be considered, as it is difficult to resource. ^89^Zr is currently the radionuclide of choice, widely used to label Abs. For clinical applications, ^89^Zr has a suitable half-life of 78 h and a low positron range, which results in optimal PET image resolution.

Several approaches for radiolabeling of nanomedicines with ^89^Zr have been explored, the majority of which use bifunctional chelators attached to the surface of the nanomaterial [11] [12]. However, this surface labeling approach using chelators has shown caveats such as modification of the chemical properties of the surface of liposomes which can lead to modified biodistribution [13].

To minimize the chemical modifications of the liposomal surface, which can lead to modified biodistribution, we propose using direct labeling using the ionophore oxine. We first described the use of ^89^Zr-oxine complex for radiolabelling and PET imaging of PEGylated liposomes[14–16]. This radiotracer has shown excellent liposome radiolabelling properties due to is lipophilic and neutral nature, allowing efficient crossing of the lipid bilayer, and its metastability results in intraliposomal radionuclide release where it binds to internal components (*i.e.* metal-binding drug molecules). This method aims to be simpler and translatable and confer a stable labeling. The two described formulations, generic Doxil and Talidox, were labeled with this approach and studied *in vitro* and *in vivo* to evaluate the label’s stability *in vivo* and usability for planned future clinical use.

## 2. Material and methods

### 2.1. Materials

Deionized water was obtained from a PURELAB® Chorus 1 Complete instrument (Veolia Water Systems LTD, UK) with 18.2 MΩ cm resistance and was used through this study. Talidox was provided by InnoMedica Holding AG (Switzerland). Doxorubicin Accord PEGylated liposomal 2mg/mL Concentrate for solution for infusion (Accord Healthcare Limited, UK) a bioequivalent of Doxil was provided by Guy’s Hospital Pharmacy (London, UK) and henceforth, will be addressed as Doxil in this work. Plain HSPC/CHOL/mPEG2000-DSPE Liposomes (50:45:5 mol/mol, 100 nm) (Formumax Scientific Inc., United States) were used as placebo liposomes (empty PEGylated liposomes). ^89^Zr was purchased from PerkinElmer as [^89^Zr]Zr-oxalate in 1.0 M oxalic acid (BV Cyclotron, VU, Amsterdam, the Netherlands). All other chemicals and reagents were commercially available and of analytical grade and used without further purification.

### 2.2. Radiolabeling feasibility

Radiolabeling feasibility experiments were performed to assess the suitability of the label *in vitro* by studying the radiochemical yield (RCY), liposome stability, and serum stability (SS) of the purified labeled liposomes. The labeling consisted of a two-step incubation radiosynthesis: 1) Production of the lipophilic ^89^Zr-oxine complex 2) Radiolabeling of the liposomes and purification.

#### 2.2.1. Production of [^89^Zr]Zr(oxinate)_4_

Labeling feasibility was carried out by preparing [^89^Zr]Zr(oxinate)_4_ using a previously described radiopharmaceutical kit [17, 18].One to 4 µL of [^89^Zr]Zr-oxalate (1M) Perkin Elmer (Netherlands) was added to 50-100 µL of the kit. The radiochemical purity of [^89^Zr][Zr(oxinate)_4_] was confirmed by radioTLC, using Whatman n°1 filter paper (Cytiva, USA) and EtOAc as mobile phase. RadioTLCs were acquired with Scan-RAM equipped with a PET probe (LabLogic Systems Limited, United Kingdom). Data was visualized with Laura radiopharmacy software (LabLogic Systems Limited, United Kingdom). Regions of interest (ROI) were drawn to quantify the area under the curve. A radiochemical purity (RCP) of oxine above 80% was deemed acceptable to continue to the second step.

#### 2.2.2. Radiolabeling of the liposomes and purification

Ten µL (1-2 MBq) of [^89^Zr][Zr(oxinate)_4_] were added to 200 µL of generic Doxil or Talidox® preheated to 50 °C. The suspension was incubated with periodical shaking for 30 min. To a previously PBS equilibrated size exclusion column (PD Minitrap™ G-25, Cytiva, USA), 500 µL of either liposomal formulation (210 µL of liposomes + 290 µL PBS) were added. The purified liposomes were eluted with 700 µL of PBS, discarding the first 200 µL, and the rest 500 µL were kept as the purified particle (*P_p_*) sample. RCY was calculated as the activity in the collected fraction divided by the *=P_p_ P_p+ R_x 100* Equation 1.

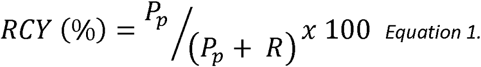

#### 2.2.3. Quality control of the radiolabeled liposomes

Quality control of the liposomes involved

#### 2.2.4. Serum stability studies

Radiochemical serum stability (SS) was tested in human serum from human male AB plasma, USA origin, sterile-filtered (Sigma-Aldrich, USA) in a ratio 1:1 incubated at 37°C for 72 h. Radiochemical purity (RP) of the liposomes was assessed using an ÄKTApurifier (GE, Sweden) equipped with a SuperoseTM 6 Increase 10/300 GL (Cytiva, USA) column running with 0.5 ml per min PBS flow. Fractions of 1 mL were collected and measured in the gamma counter 1282 COMPUGAMMA CS (Wallac, Finland). The activity of the fractions with the same retention time as the unlabeled liposomes (*P_p_*) was divided by the activity of the total fraction (*A_T_*) and represented *=P_p_ P_p+ AT_x 100* Equation 2 *SS %=P_p_ P_p+ AT_x 100*.

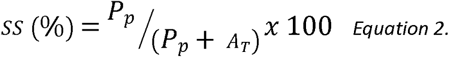

### 2.3. Preclinical production of ^89^Zr-labeled liposomes

To achieve the required activity concentration for preclinical studies, [^89^Zr]Zr-oxalate underwent a conversion/concentration step. Forty-eight MBq were diluted in 250 µL of Chelex-treated water and loaded in a pre-conditioned Sep-Pak Accell Plus QMA Plus Light Cartridge (Waters, USA), the activity was eluted with 500 µL of HCl in fractions, the fractions with most activity were joined (totaling 27 MBq) and dried under a N_2_ stream at 50 °C (for 20-30 min), after that the solid remaining was incubated with 80 µL the radiopharmaceutical oxine kit for 15 min at room temperature [17]. Twenty-five µL of the kit was added to Doxil (500 µL) and 15 µL to Talidox (500 µL).

The radiochemical purity of the injected liposomes was tested with Amicon® Ultra 0.5 mL 30 K filters (Millipore, Merk, Germany).10 µL of the liposome solution was added to 400 µL of PBS to the Amicon® filter followed by washing three times with 400 µL of PBS, measuring the filtrate and the remaining activity in the liposomes. If the RCP was below 95%, the liposomes underwent a purification step by washing three times in PBS in an Amicon® Ultra 0.5 mL 30 K filter and tested again for radiochemical purity. DLS size measurements were performed. The final concentration of Doxorubicin was 0.125 mg per dose for both formulations. The lipid concentration of Doxil was adjusted with Placebo liposomes to the same concentration as Talidox (3.2 µmol per mouse).

### 2.4. Density functional theory

The intraliposomal coordination chemistry and high stability of ^89^Zr radiolabeling was studied with density functional theory (DFT). DFT was used for all calculations with the Gaussian 16 program (version C.01) [19]. Structures were geometry optimized without symmetry constraints, in an aqueous solution using the SMD self-consistent reaction fields [20]. Each optimized structure was confirmed as local minima on the PES via frequency calculations. DFT calculations were undertaken with the B3LYP, M06-2X, ωB97xd, or PW6B95D3 functional, and either 6-31G(d,p), LANL2DZ, LANL2TZ, DGDZVP, Def2-TZVP, or Sapporo-TZP-2012 basis sets (sourced from the online Basis Set Exchange) [21–29]. Mixed basis sets were used via the ‘gen’ keyword, with the Sapporo-TZP-2012 basis set for Zr and 6-31G(d,p) for ligand atoms. When first-order SCF failed to converge, quadratically convergent SCF was employed F(via the ’scf = xqc’ keyword). Default values for temperature and pressure were used (298.15 K, 1 atm). The default integration grid (UltraFine, 99 radial shells, 590 angular points per shell) was used throughout. Root mean square error (RMSE), mean absolute error (MAE), and continuous shape measurement (CShM) were chosen to analyze the optimized geometries [30].

Doxorubicin was fragmented to retain most of its electronic system relevant for coordination to reduce the computational cost of calculation, this fragment was referred to as Dox in the computational study (Supplemental Figure S1), oxinate as Ox, and their protonated counterparts as HDox and HOx respectively. As suggested by Kheirolomoom and coworkers, Dox was coordinated to zirconium via its least sterically hindered aldol [31]. The converged geometry from Gaussian was used as the only conformer for each structure under study, whilst multiple isomers for each complex were calculated and the lowest energy isomer taken forward. Boltzmann population distribution was used to estimate the distribution of zirconium species within the liposome at different simulated pHs.

An appropriate basis set and functional was determined via benchmarking their geometries against a crystal structure of Zr(oxinate) [29] [32] which would be the left-most complex on the reaction coordinate for the DFT experiments. The chosen basis sets were judged employed with the functional B3LYP (Supplemental TABLE S1). The best performing basis sets were Def2-TZVP and Sapporo-TZP-2012/6-31G(d,p), which calculated geometries with the lowest MAE and RMSE; Sapporo-TZP-2012/6-31G(d,p) was chosen for its faster optimization time in calculating the Zr(oxinate) geometry. This basis set was then used in benchmarking the chosen functionals (Supplemental TABLE S2). ωB97xd and M06-2X scored almost identically in the error metrics, CShM, and relative optimization time. M06-2X was taken forward as a better match for the chosen self-consistent reaction field (SCRF), SMD, an SCRF that was parameterized using other Minnesota functionals (M05-2X),2 and as such, the expectation was that M06-2X would perform better in calculating accurate Gibbs energy values compared to ωB97xd.

The pH range inside the aqueous environment of the liposome throughout the experiment is estimated to fall between neutral and acidic (intraliposomal pH 5 (NH_4_)_2_SO_4_ buffer) [33]. Therefore, the experiment and reaction schemes were designed under a simulated neutral or acidic environment. This required identifying the possible ligands that exist in free solution under different pH conditions and investigating the equilibrium of proton transfer between HDox, Dox, HOx, and Ox (Supplemental Scheme S1). This suggests that under neutral conditions (Supplemental Scheme S2), free Ox will be consumed by any free HDox (ΔG = -27 kJ mol^-1^), liberating free Dox for complexation. Under acidic conditions (Supplemental Scheme S2), Ox and Dox will both be protonated to HOx and HDox (-67 and -40 kJ mol^-1^ respectively). These results match the predicted pKa values from Ji and co-workers’ graph-convolutional neural network pKa predictor [34] – 7.8 for HOx, 7.1 and 7.2 for HDox.

### 2.5. *In vivo* model in BALB/c mice bearing 4T1 orthotopic tumors

#### 2.5.1. Cell culture and inoculation

4T1 cells were cultured in complete RPMI medium supplemented with L-glutamine (2 mM), Penicillin (100 units)/Streptomycin (0.1 mg) per mL, and fetal bovine serum at 37 °C in a humidified atmosphere with 5% CO_2_. The cells were detached using TrypLE Express Enzyme ™ (Gibco, Thermo Fisher, USA) and washed with 50 ml PBS twice by centrifugation at 500 *g*. The cell suspension was concentrated to 10 million cells per ml of PBS.

#### 2.5.2. *In vivo* imaging

Animal imaging studies were ethically reviewed and carried out in accordance with the Animals (Scientific Procedures) Act 1986 (ASPA) UK Home Office regulations governing animal experimentation. One million of 4T1 cells in 0.1 ml of PBS were subcutaneously implanted in the first mammary pad of BALB/c female mice aged 56-60 weeks (Charles Rivers, UK).

After 1 week of tumor growth, the mice were anesthetized using gaseous isoflurane with 1 L per min O_2_ flow. The mice body temperature was regulated at 37 °C using a heating block. Mice were injected with either approximately 0.8 MBq of [^89^Zr]Zr-Doxil, 0.7 MBq of [^89^Zr]Zr-Talidox®, or PBS intravenously through the tail vein (n=4 for imaging groups, n=2 for controls).

The mice were transferred to the nanoPET/CT scanner (Mediso, Hungary) with acquisition software Nucline-2.01.020 (Mediso, Hungary). All animals were imaged for 30 min followed by a CT scan. The imaging was repeated at time points 24 h, 48 h, and 72 h, with the duration of scans corrected for decay. Tail vein blood withdrawals were collected in 10 µL capillary tubes at regular intervals and weighed to determine blood pharmacokinetics. Static PET reconstruction was performed using standard settings on Nucline v.0.21 software.

#### 2.5.3. Image analysis

Images were analyzed and preprocessed using VivoQuant software (version 3.5, InviCRO) to show the image scales in %IA/g (percentage Injected Activity per gram of tissue) and the bed and probe was removed to allow Maximum Intensity Projections (MIPs) visualization. For the ROIs analysis the images were preprocessed to Bq and ROIs were drawn on the organs of interest for each time point. The resulting Bq per ROI was divided by the injected dose and voxel volume. The resulting %IA/mL values were plotted against time (0 to 72 h) and AUC analysis was performed using GraphPad 10.2.

#### 2.5.4. *Ex vivo* biodistribution and blood pharmacokinetics

At the end of the last imaging session (72 h) the mice were culled by cervical dislocation and the organs were harvested for biodistribution measurements. Each organ was weighed and measured in the gamma counter, together with standards prepared from a sample of the injected material, to calculate the %IA/g values for each tissue sample. The %IA/g for blood was also calculated from blood samples, plotted against time and the data fitted to a single exponential decay model to calculate the biological blood circulation half-life of each liposome.

#### 2.5.5. Autoradiography

The tumors were snap-frozen in isopropanol at -80 °C and sectioned to 10 µm thickness to perform autoradiography. Half of the tumors were kept at -80 °C for CFT. The sections were exposed to GE phosphor plate overnight. The phosphor plate was developed with the Phosphoimager tool of Typhoon Amersham (GE, USA). Images were processed in the Imaging software ImageQuantTL 10.0 -261 (Cytiva, USA).

#### 2.5.6. Cryofluorescence tomography (CFT)

CFT was performed on tumors using the Xerra CFT cryomacrotome (Xerra™, Emit Imaging, Baltimore, MD). The CFT uses a 12-megapixel camera, 6 channel laser module and 7 interchangeable emission filters to automatically slice and sequentially image, providing isotropic images in white light and fluorescence. Tumors were prepared as follows.

The -80 °C frozen tissues previously described were embedded in optimal cutting temperature (OCT) compound in a block measuring 8 x 6 x 4 cm and frozen at -20 °C. For CFT, the frozen block was transferred to the cutting stage inside the cryo-chamber at -14 °C and was serially sectioned at 20 µm increments. The block face was imaged at each sectioning plane in white light (WL) and fluorescence imaging (DOX: ex 470 nm/em 620 nm) using Xerra Controller software (Emit Imaging). Raw WL and fluorescent images were reconstructed in the Xerra Reconstruction software (Emit Imaging) and MHD files (MetaImage Metaheader files) of the whole block were generated for WL and fluorescent images. Maximum intensity projections (MIPs) of the whole block fluorescent images were then created using ImageJ software.

### 2.6. Statistics

Descriptive data are presented as mean ± standard deviation unless otherwise mentioned. Results were considered statistically significant at *p* < 0.05, without correction for multiple comparisons. Comparison of groups, non-linear regression analysis and generation of graphs was performed using GraphPad Prism version 10 (GraphPad Software 10.4).

## 3. Results and Discussion

### 3.1. Radiolabeling feasibility studies

Radiochemical yield for both formulations in feasibility studies was 59 ± 5 % (n=4) for Talidox® and 62 ± 8 % (n=4) for generic Doxil. The radiochemical stability in human serum after 72 h incubation at 37°C was 77 ± 11% (n=3) for Talidox® and 76 ± 16 % (n=3) for Doxil (Figure 1A). The stability is comparable to previously imaged [^89^Zr]Zr-labeled antibodies (Ab) [35], which was suitable for preclinical imaging. Although [^89^Zr]Zr−DFO labeled Ab have generally shown superior *in vitro* serum stability values (<95%), however, it does not directly translate to *in vivo* stability in all cases, and even showing indirect correlation in some instances [36]. Therefore, stability was deemed sufficient to progress into *in vivo* studies. Hydrodynamic size (Z-average), polydispersity index (PDI) and surface charge (zeta-potential) of the liposomal formulations before and after labeling were not significantly different, as shown in **Error! Reference source not found.**B-D. Minimal impact on hydrodynamic size confirmed no modifications to the size of the liposomes, an important property defining the circulation half-life as well as biodistribution. The polydispersity index remained below <0.3 after radiolabeling, which is considered acceptable for clinical formulations of phospholipid-based nanocarriers indicating homogenous populations of liposomes. Finally, surface charge also remained similar, suggesting that the membrane has not been modified and thereby limiting any impact of biodistribution and pharmacokinetics.

**Figure 1.**
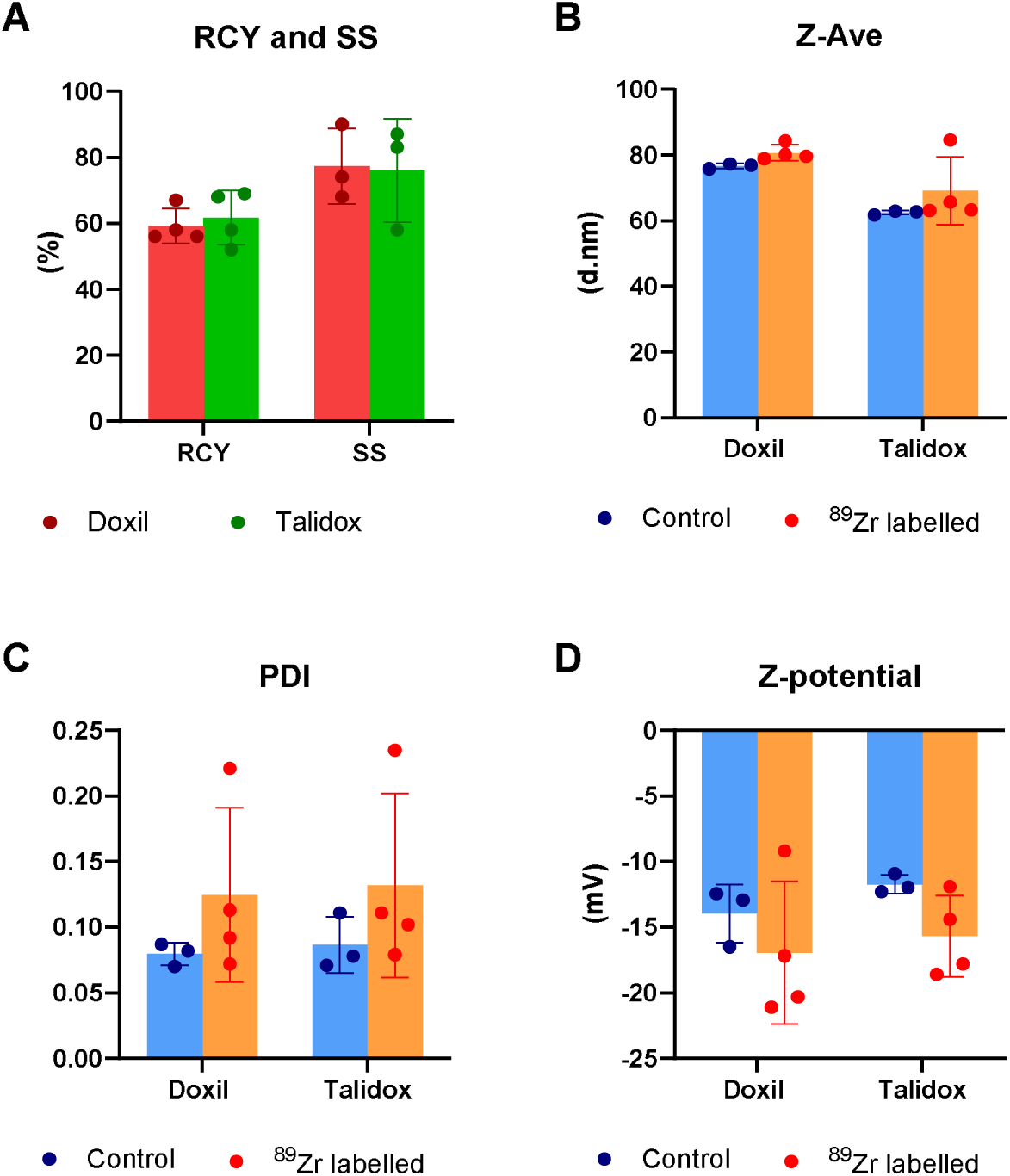
A) Comparison of the radiochemical yield (RCY) and serum stability (SS) at 72 h for generic Doxil (Red) and Talidox (Green). B) Z-average (Z-Ave) in nm of Talidox and Doxil before (blue) and after (orange) labelling. C) Polydispersity index (PDI). D) Z-potential in mV. All column heights represent mean values and error bars represent standard deviations.

This radiolabeling of liposomal formulations would be compatible with GMP standards. The typical clinical patient dose of Doxil for an average woman (1.6 m^2^ body surface area) is 16 – 40 mL depending on the pathology [1] and trials with Talidox have used *ca*. 64 mL also considering an average woman [37]. This would allow a minimum addition of up 800 µL of oxine kit (5 % of the total labelling volume of 16 mL) which effectively permits addition of 15 % of [^89^Zr]Zr-oxalate (120 µL of [^89^Zr]Zr–oxalate, >120 MBq) leading to clinical dose of >40 MBq of purified liposomes (under the assumption of minimal RCY of 30%). This dose of radiolabeled liposomes is feasible for conventional PET/CT studies [38]. Moreover, with the advent of clinical total-body PET imaging scanners, this required dose could likely be reduced due to the higher sensitivity of these novel instruments [39]. This is in contrast with other reported methods, which include an extensive synthesis scheme requiring solvents such as chloroform only approved as impurities (EMEA exposure limits of 0.6 mg dose/60 ppm)[40, 41]. Such residual impurities are often measured by gas chromatography [42] which is often cumbersome and sparsely available in traditional radiopharmacies. The ^89^Zr-oxine labelling kit was optimized and designed to be similar to the already established [^111^In]In-oxine labeling kit, which should facilitate regulatory approval for clinical studies/trials [43].

Thereby, this labeling fulfilled the two objectives of feasibility studies. Firstly, the labeling process was simple, highly reproducible, and easily scalable to clinical doses. Secondly, the resulting radiolabel was stable and did not modify the physicochemical properties of the liposomes.

### 3.2. Preclinical production of ^89^Zr-labelled liposomes

To achieve the high specific activity required for preclinical studies, the radiolabelling process was modified. In the modified radiolabelling process, pre-processing of ^89^Zr was performed to achieve higher specific activity. This involved loading of the [^89^Zr]Zr-oxalate on a strong hydrophilic and anionic exchanger cartridge and eluting concentrated [^89^Zr]ZrCl_4_ in a small volume. This concentrated ^89^Zr was then used for the radiolabelling of liposome formulations. The radiochemical yield of Doxil was 30.3% with a RCP of 92.4% thereby, needing a purification step as the RCP was below the QC limit of 95%). The purification was performed using Amicon® Ultra Centrifugal Filters providing radiolabelled Doxil with an RCP of 97.4% and accounting for the losses compared to feasibility studies. Talidox had a decay-corrected radiochemical yield of 48% with RCP of 98.5% and thereby, requiring no further purification. As expected, no physicochemical impact of radiolabelling was observed on hydrodynamic size, surface charge, and polydispersity for both Doxil and Talidox, comparable to the unlabelled liposomes.

The need for high specific activity was necessary for the preclinical step due to the limits on volume administered intravenously and liposomal doxorubicin dose in mice (maximum limits of administration: 200 µL *i.v.* for the ∼20 g mice used in this study and 5 mg/Kg of doxorubicin). However, it is important to note that the volume administration limits for patients in a clinical setting are much larger [1] and thereby, removing the need of the concentration step, as discussed in the previous section.

### 3.3. DFT calculations support the experimentally observed high ^89^Zr radiolabelling stability of doxorubicin-loaded liposomes

Density functional theory was used to model the reaction between Zr(oxinate)_4_ and doxorubicin in an aqueous solution to replicate the reaction conditions within a liposome. Experiments were conducted in simulated acidic and neutral environments at the M06-2X/SAPPORO-TZP-2012/6-31G(d,p) level. Geometry optimization and frequency calculations provided Gibbs energy values for each arrangement of the ligand substitution between Zr(Ox)_4_ and Dox (a fragment of doxorubicin) via a dissociative mechanism (Figure 2). In both acidic and neutral environments, the substitution was shown to be thermodynamically feasible with Boltzmann distribution calculations suggesting that the neutral scenario would have a majority of Zr(Dox)_4_ (99.7%). The acidic scenario would have a majority of [Zr(Ox)_1_(Dox)_2_]^+^ (97.2%). Please refer to supporting information for in-depth simulation results for the neutral and acidic environment (Supplemental Scheme S1 and S2).

**Figure 2.**
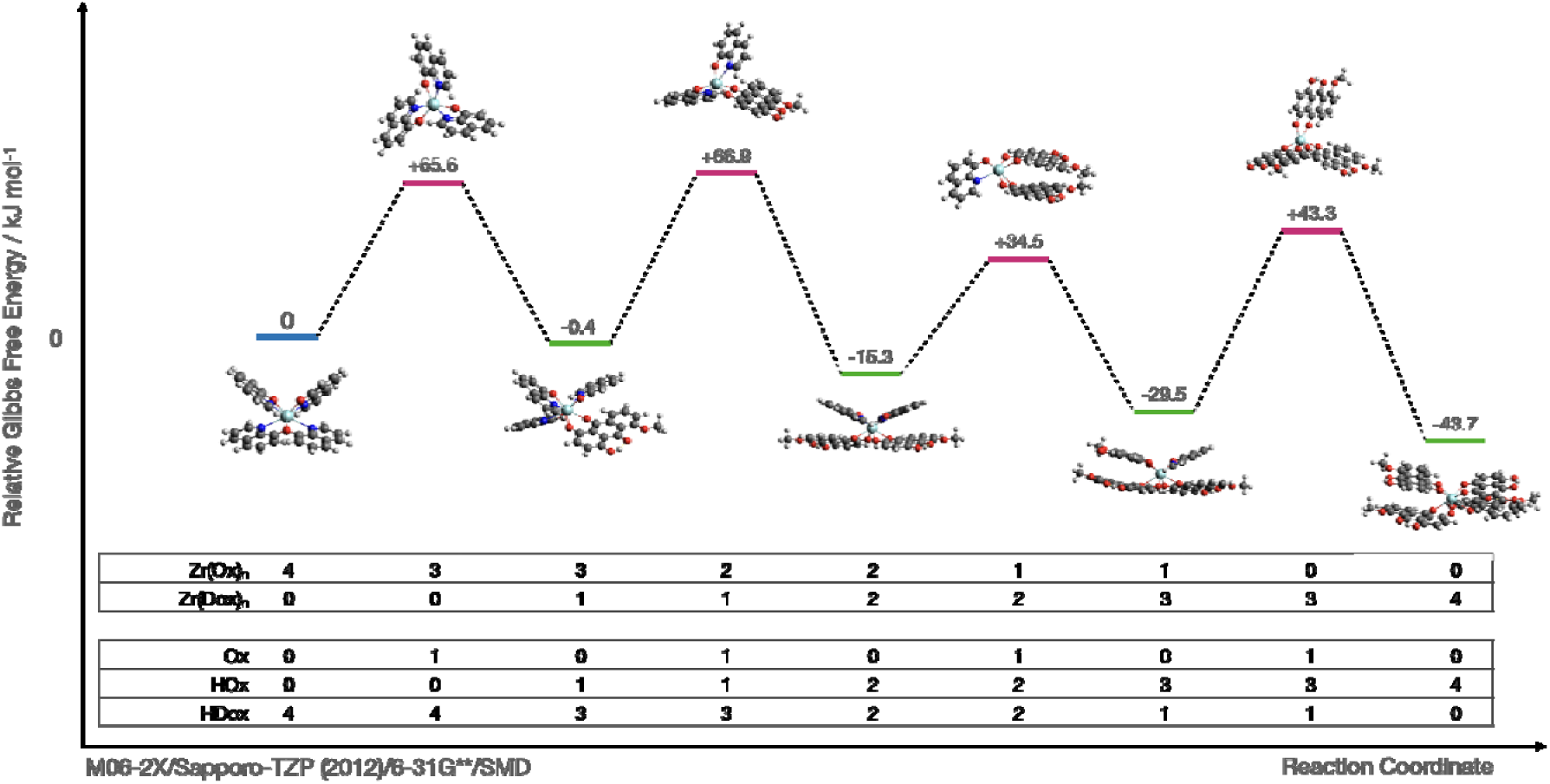
Calculated Gibbs energy levels and accompanying relative change in Gibbs energy in kJ mol-1 relative to the starting arrangement of Zr(Ox)_4_ and four HDox ligands in aqueous solution, in the conversion of Zr(Ox)_4_ into Zr(Dox)_4_ under simulated neutral conditions, at M06-2X/Sapporo-TZP-2012/6-31G(d,p)/SMD level. Upper table inset: count of Ox or Dox coordinated to zirconium in that arrangement; lower table inset: count of uncoordinated ligands Ox, HOx, and HDox in the aqueous solution to balance stoichiometric equivalence between arrangements.

### 3.4. *In vivo* imaging, biodistribution, and pharmacokinetic comparison of [^89^Zr]Zr-Doxil and [^89^Zr]Zr-Talidox

After successful experimental and computational confirmation of the stability of radiolabelled liposomal formulations, the tracking of radiolabelled Doxil and Talidox was performed in a syngeneic breast cancer 4T1 model in immunocompetent BALB/c mice. The 4T1 tumour model was chosen due to its spontaneous, metastatic tumour growth in BALB/c mice and that closely mimics late-stage human breast cancer. The tumour cells were implanted on the first mammary pad to avoid any tumour accumulated radiotracer signal interference with PET signal emanating from expected liver and spleen accumulation of liposomal nanomedicines.

^89^Zr labelling allowed tracking of Doxil and Talidox for up to 72 h due to its long half-life and high stability of the radiolabel. Representative images of the biodistribution of [^89^Zr]Zr-Doxil and [^89^Zr]Zr-Talidox observed at t = 20 min, 24 h, 48 h and 72 h are shown in Figure 3. At early timepoints of t = 20 min and 24 h, both liposomal formulations were observed in high percentage in blood circulation. After 48 h, the majority of the accumulation was observed in the reticuloendothelial system (RES) organs *i.e.* spleen, liver, as well as in the tumour. At 72 h, this effect was even more pronounced with further increased accumulation in the spleen, liver, and tumour. These observed results are in line with similar PEGylated liposomal formulations [44].

**Figure 3.**
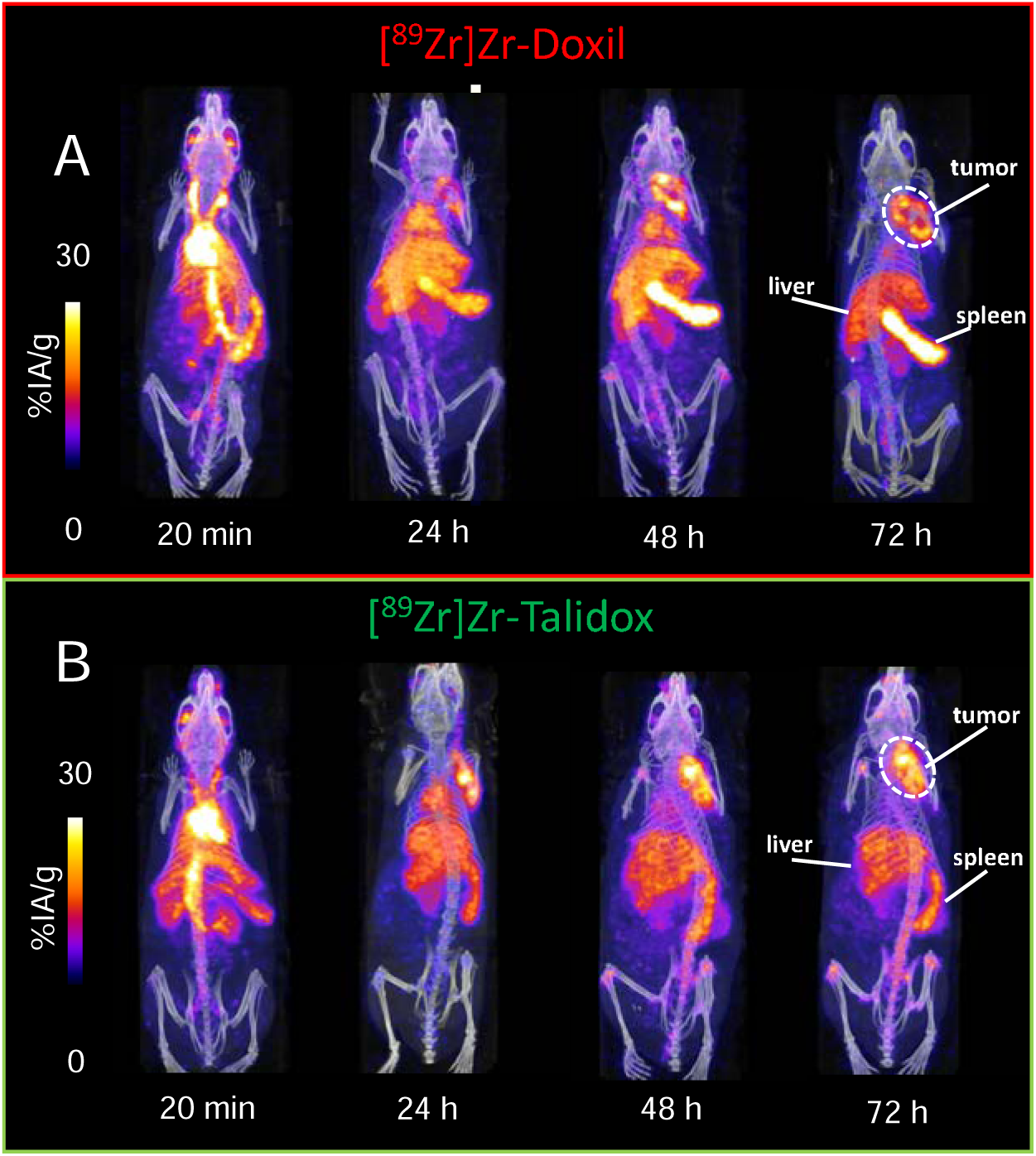
Representative PET/CT images of one mouse per group. A) Doxil injected mouse at 20 min, 24 h, 48 h, and 72h, showing progressive accumulation in tumour, liver and spleen. B) Talidox injected mouse at 20 min, 24 h, 48 h, and 72h, showing progressive accumulation in tumour, liver and spleen.

Quantitative image analysis – region of interest (ROI) analysis – also showed similar profiles for both formulations, as shown in Figure 4A,B. However, differences in the areas under the curve (AUCs) showing cumulative accumulation of liposomal formulations were significant for the heart/blood, bone, kidney, liver, lungs and very pronounced in spleen (see Figure 4C). A higher spleen accumulation for the Doxil formulation was observed as expected due to the bigger size of Doxil [45]. However, the increased accumulation of Doxil in the spleen did not lead to substantial decreased tumour delivery of Doxil compared to Talidox. This could be explained by the faster clearance of Talidox by the liver, which could further explain the higher observed bone uptake of Talidox due to the loss of free ^89^Zr label.

**Figure 4.**
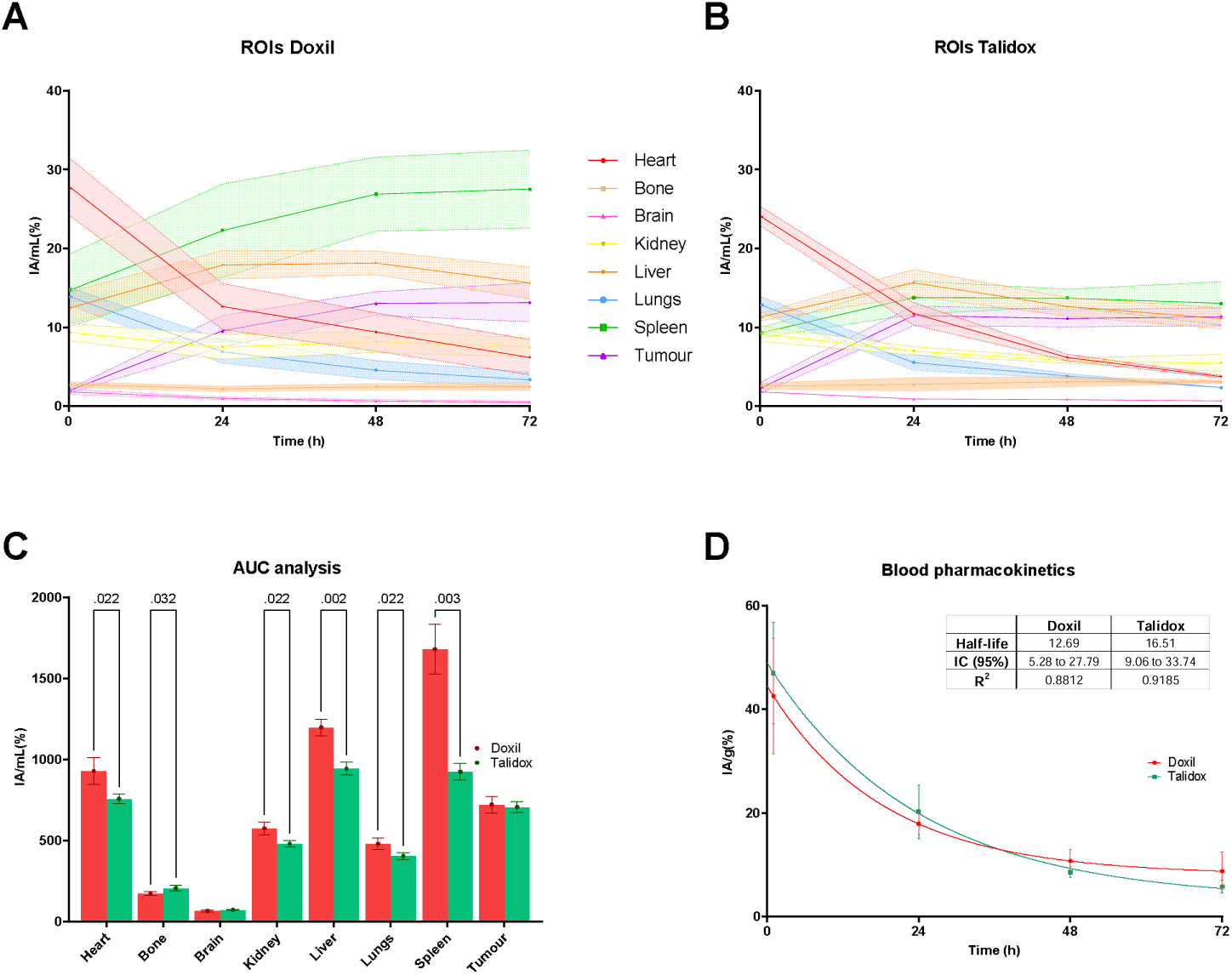
A) ROI analysis of ^89^Zr-Doxil mice of organs of interest. The solid line connects the calculated means in each time point and the shadowed area the standard deviation. B) ROI analysis of ^89^Zr-Talidox mice of organs of interest. C) Comparison of ^89^Zr-Doxil and ^89^Zr-Talidox AUCs_(0-72 h)_ of organs of interest. The numbers represent the p value where a significant difference is found between formulations. D) Blood pharmacokinetics graph with the values of %IA/g versus time. A single exponential decay fitted and represented as a solid line.

The pharmacokinetic analysis shown in Figure 4D showed a statistically significant longer blood half-life for [^89^Zr]Zr-Talidox 16.5 h (R^2^ = 0.92) than for [^89^Zr]Zr-Doxil 12.7 h (R^2^ = 0.81), but the %IA/g for the blood compartment in the final timepoint of 72 h was not significantly different (Figure 5 5). This is in contrast to the half-life reported in other publications for Doxil, in the range of 20-30 h [44]. Differences in the results could be due to using a different kinetic model for fitting the blood clearance. We chose a simple exponential one phase model because it gave similar R^2^ to more complex models as the two-phase model used in previous studies [46]. With our reduced sample size (n=4) and their variability in the first part of the curve, the two-phase model could not resolve the first phase.

**Figure 5.**
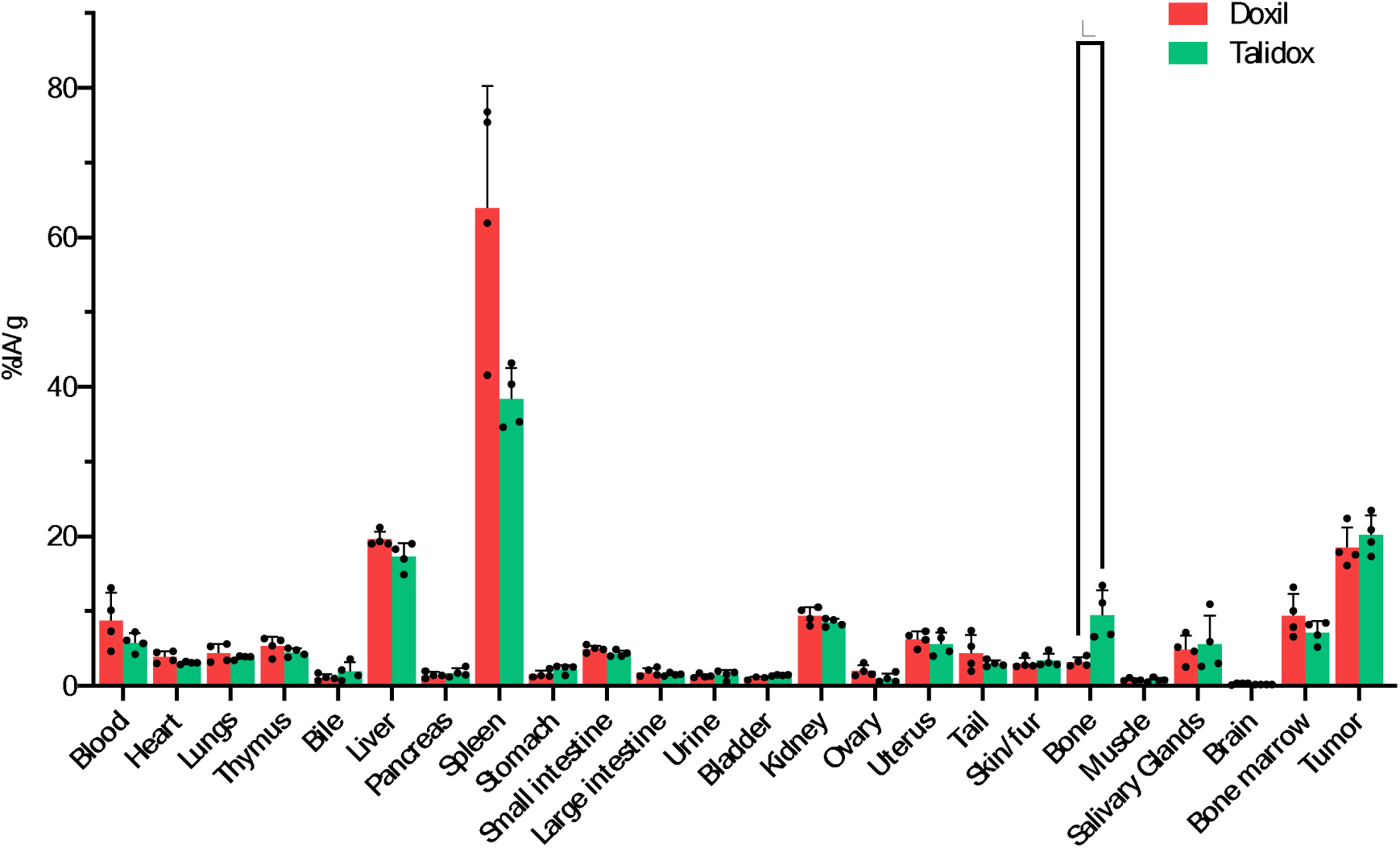
5.Biodistribution of Doxil (red) and Talidox (green) at 72h post adminstration. The %IA/g is presented for each organ, and the numerical results are the mean of each group the error bar represents the standard deviation of the mean.* P ≤ 0.05

In the context of biodistribution of liposomal formulations in the bones, it is important to note that free ^89^Zr has been previously known to accumulate in the hydroxyapatite matrix of the bone [47]. Due to the limited spatial resolution of PET imaging and the abovementioned accumulation of free ^89^Zr in bone mineral, image analysis using ROIs cannot be used to determine the PET signal originating from liposomes that might accumulate in the bone marrow. This issue was resolved in the *ex vivo* biodistribution studies by separating the bone marrow from the bone by high-speed centrifugation (Figure 5, Table S3). The radioactive signal in bone mineral measured using the gamma counter was as 3.2 ± 0.64 % (n=4) IA/g for Doxil and 9.5 ± 3.3 % (n=4) IA/g for Talidox, which was significantly different (P value = 0.0286). In contrast, the uptake in the bone marrow was higher for Doxil (9.4 ± 2.9 %; n=4) *vs*. Talidox (7.2 ± 1.5 %; n=4), but the difference was not statistically significant. The higher observed bone uptake for Talidox is likely due to a higher clearance of Talidox by the liver, as the bone uptake was more pronounced in the later time points (Figure 3). This observation, however, is unlikely to be due to the instability of the radiolabelling as that would lead to high bone uptake in the initial 24 h post-administration [48].

### 3.5. Autoradiography and cryofluorescence tomography (CFT) inform intratumoural biodistribution

There was high uptake in the tumour for both formulations as visualized qualitatively and quantitatively by PET imaging and *ex vivo* biodistribution studies ([^89^Zr]Zr-Doxil = 18.5 ± 2.4 % IA/g (n=4), [^89^Zr]Zr-Talidox = 20.2 ± 2.3 % IA/g (n=4)) but did not inform about intratumoural biodistribution and heterogeneity. We explored intratumoural biodistribution qualitatively through autoradiography of thin tumour sections (10 µm). The observed distribution within each section hinted towards a more even distribution of [^89^Zr]Zr-Talidox (Figure 66B) in the core compared to [^89^Zr]Zr-Doxil (Figure 6A), which remained near the main blood vessels on the periphery of the tumour. This observation supports previous reports that Talidox has better penetration in solid tumours [45], and positively impact the outcome of the therapy in solid tumours.

**Figure 6.**
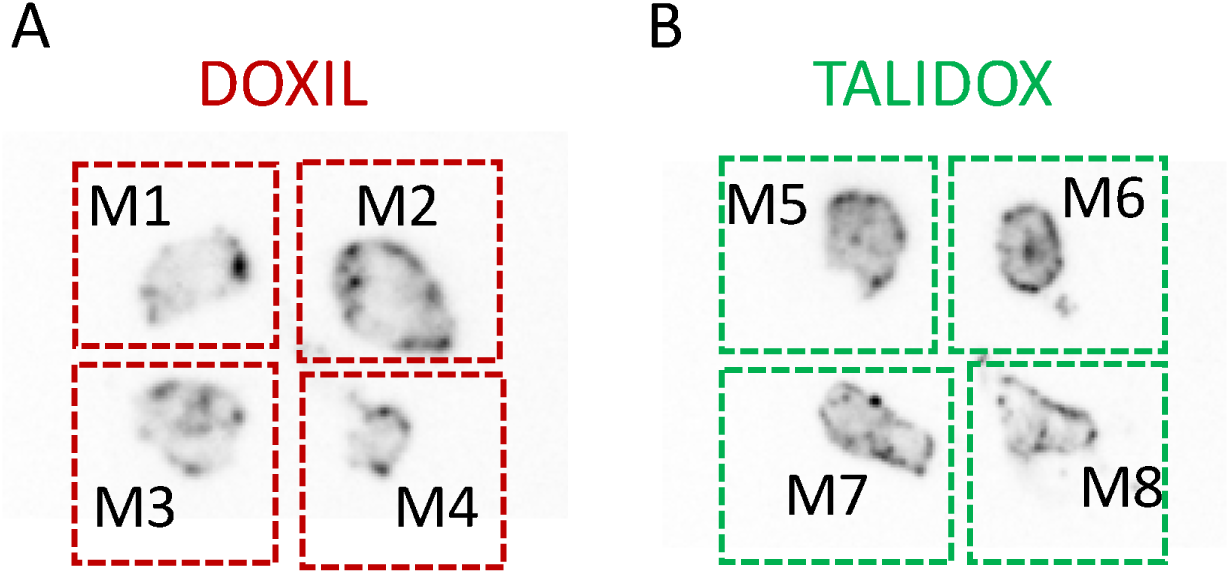
Autoradiography of sections of all tumors. A) contains a slice of 10 µm of each mouse tumor treated with [^89^Zr]Zr-Doxil and B) treated with [^89^Zr]Zr-Talidox.

Cryofluorescence tomography (CFT), a whole tissue fluorescence imaging modality, allowed for the visualization of the accumulation of Doxorubicin loaded within both liposomal formulations. **Error! Reference source not found.** shows the MIP (maximum intensity projection) maps created from CFT readings, displaying the fluorescence signal across all tumour samples in the Doxorubicin energy wavelengths. Although the non-treated tumours (negative controls: M9 and M10) exhibited some degree of tissue autofluorescence, the doxorubicin-treated tumours (M1–8) consistently showed qualitatively higher fluorescence intensities. These results suggested that liposomal treatment delivers the doxorubicin to the tumours, consistent with the PET findings above.

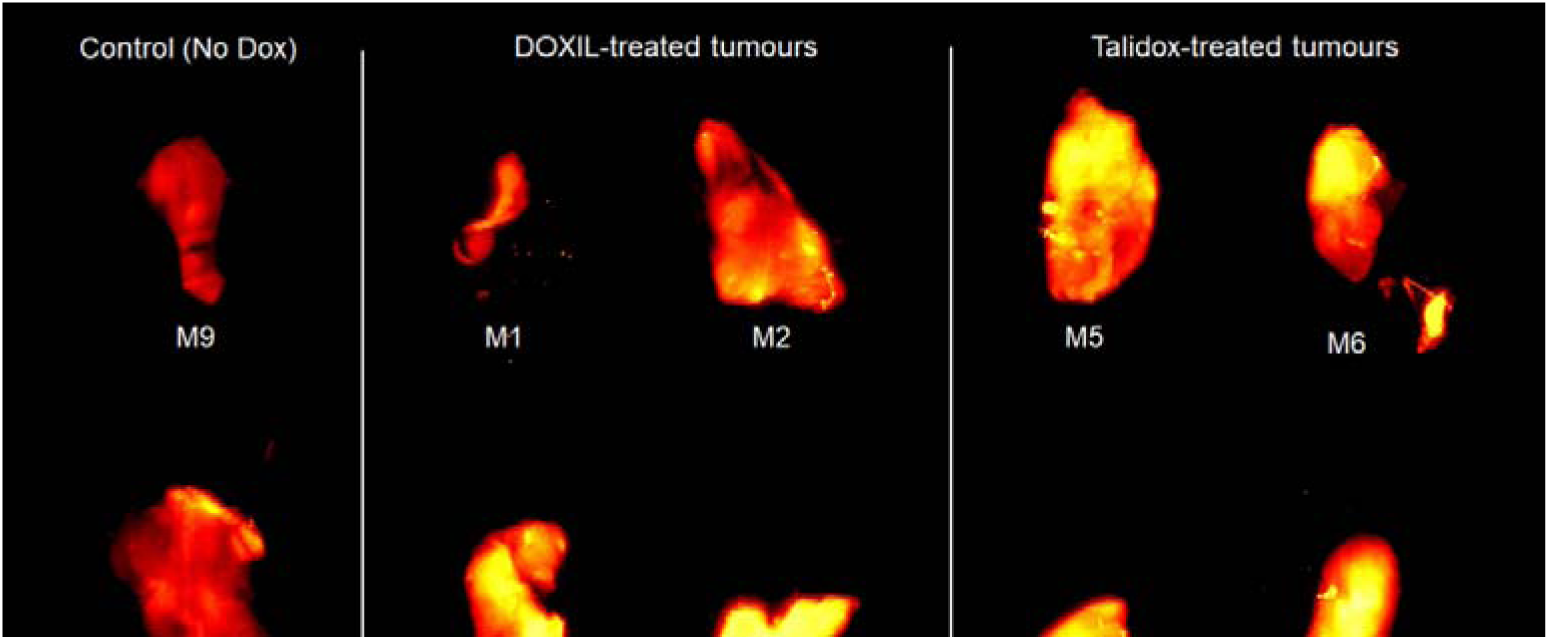

We must consider that the study had limitations. A relatively small sample size (n = 4) was used for the *in vivo* studies and increasing this may have resulted in statistically significant differences in some organs (*e.g.* spleen). Furthermore, the CFT tumour imaging data showed high tissue autofluorescence signal, which is a result of the suboptimal excitation/emission properties of doxorubicin as a fluorophore for tissue imaging.

## Conclusions

Liposomal formulations present advantages to traditional chemotherapy, including reduced side effects and similar effectiveness in solid tumours with properties that lead to accumulation via the EPR effect. Nevertheless, there is currently no tool available to identify these tumours and stratify patients for liposomal nanomedicine treatment, which leads to limited indications and potentially patients missing a more tolerable chemotherapy.

The radiolabelling methodology proposed here would allow to use PET to image and stratify patients from the first dose, study the effect of coadjutant therapy (*e.g.* radiotherapy) or identify therapy resistance. The radiolabelling has been designed to produce minimal changes in the liposomes and would be easy to translate meeting GMP requirements. This approach can also be used as a de-risking strategy in the preclinical stages to test and identify novel formulations to progress through expensive clinical trials.

Also importantly, the direct labelling of doxorubicin containing liposomes has been proved stable *in vivo* and *in vitro*, and for the first time we present DFT data that support the experimental findings, with evidence that ^89^Zr binds strongly to intraliposomal doxorubicin.

## Acknowledgements

This work was supported by the Centre of Excellence in Medical Engineering funded by the Wellcome Trust and the Engineering and Physical Sciences Research Council (EPSRC) (grant number WT 203148/Z/16/Z); EPSRC Programme Grant (EP/S032789/1 ‘‘MITHRAS’’). PET scanning equipment at KCL was funded by an equipment grant from the Wellcome Trust under grant no WT 084052/Z/07/Z. Radioanalytical equipment was funded by a Wellcome Trust Multiuser Equipment grant: a multiuser radioanalytical facility for molecular imaging and radionuclide therapy research. The authors acknowledge support from the National Institute for Health Research (NIHR) Biomedical Research Centre based at Guy’s and St Thomas’ NHS Foundation Trust and KCL (grant no IS-BRC-1215-20006). The authors acknowledge the support of InnoMedica Holding AG for providing Talidox liposomes used in this study. The authors thank Dr Filipa Mota (Perspective Inc.) for facilitating access and running samples on the cryo-fluorescence Tomography (CFT) equipment based at Barts Cancer Institute, QMUL. The authors would like to thank the technical and preclinical team members of the Imaging Chemistry and Biology Department in the School of Biomedical Engineering and Imaging Sciences.

---

**Figure S1.**
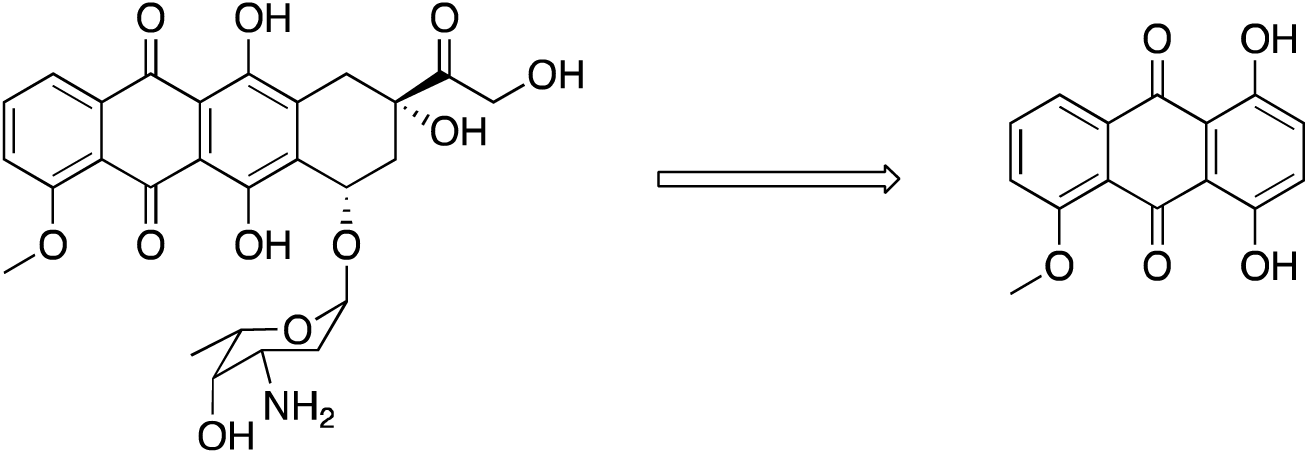
Fragment of doxorubicin used in the DFT study to provide the ‘HDox’ fragment.

**Scheme S1.**
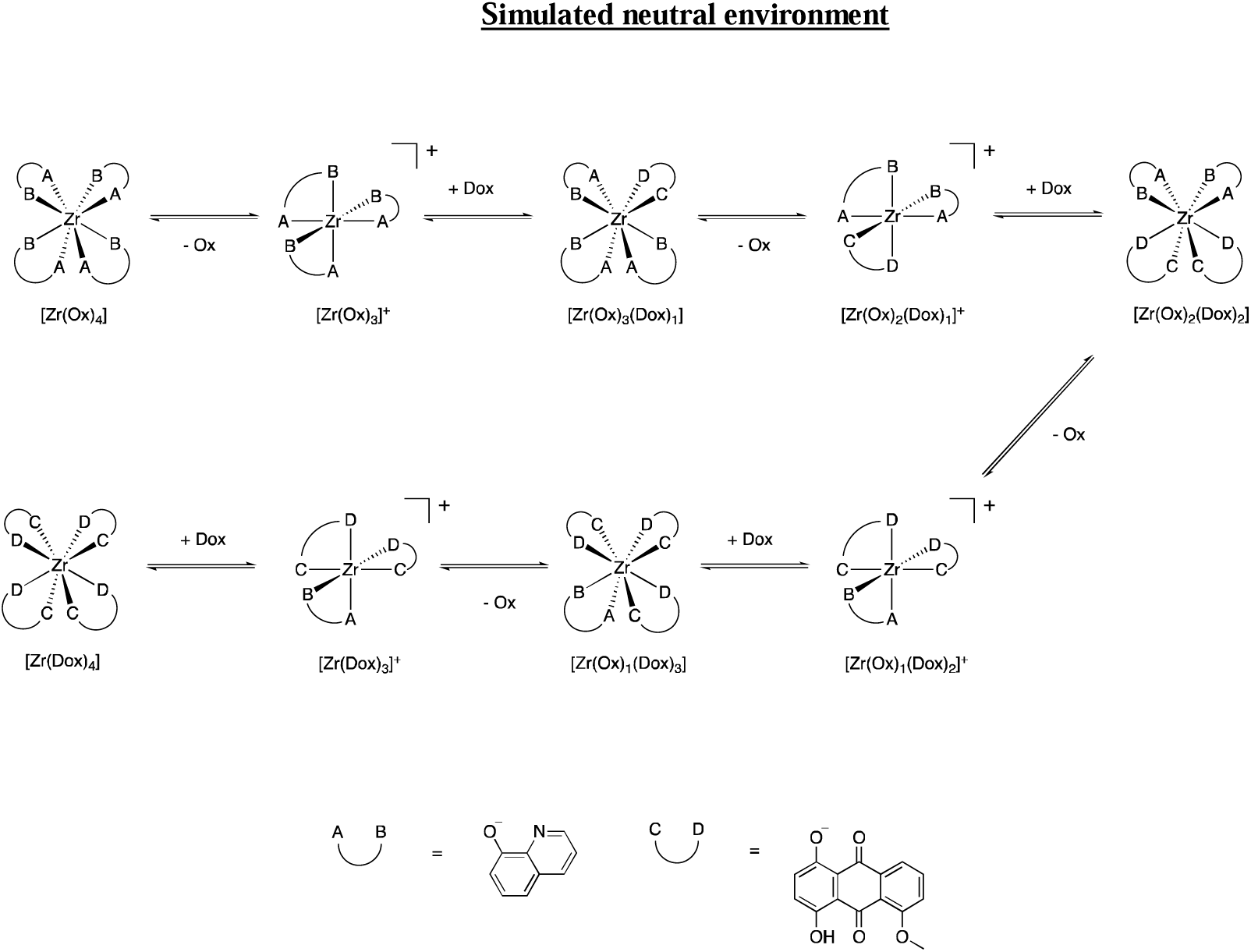
Reaction scheme for the conversion of Zr(Ox)_4_ to Zr(Dox)_4_ via a dissociative pathway in a simulated neutral environment; proton exchange between Ox and HDox [[Scheme 1, Reaction 1]], provides free Dox for complexation.

**Scheme S2.**
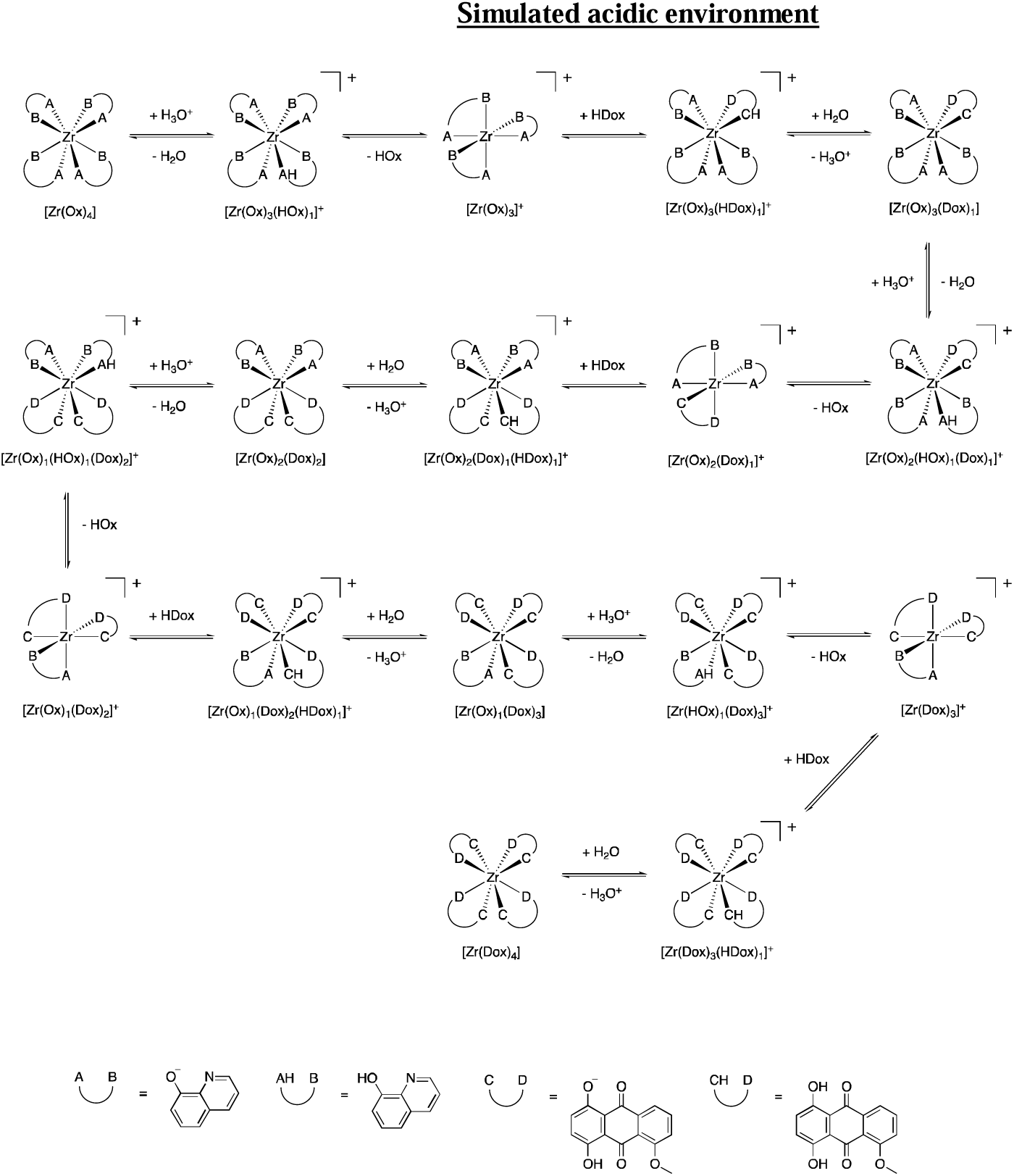
Acidic environment reaction scheme of Zr(Ox)_4_ conversion to Zr(Dox)_4_ *via* a dissociative pathway.

**TABLE S1.** Comparison of error parameters for the DFT optimised geometries of Zr(oxinate)_4_ in aqueous solution for different basis sets and the B3LYP functional.

| Basis Set<sup>a</sup> | Relative optimisation time | MAE (all)<sup>b</sup>/Å | MAE (Zr-Donor)/Å | MAE (Zr-N)/Å | MAE (Zr-O)/Å | RMSE/Å | CShM |
| --- | --- | --- | --- | --- | --- | --- | --- |
| LANL2DZ | 1.0 | 0.135 | 0.144 | 0.113 | 0.169 | 0.534 | 1.755 |
| LANL2TZ | 1.4 | 0.115 | 0.194 | 0.226 | 0.155 | 0.474 | 1.499 |
| DGDZVP | 2.2 | 0.121 | 0.202 | 0.218 | 0.185 | 0.507 | 1.701 |
| Def2-TZVP | 15.4 | 0.106 | 0.182 | 0.241 | 0.091 | 0.470 | 1.626 |
| Sapporo-TZP-2012/6-31G(d,p) | 2.1 | 0.112 | 0.182 | 0.242 | 0.086 | 0.465 | 1.607 |
| Experimental XRD<sup>c</sup> |  |  |  |  |  |  | 1.337 |

<sup>a</sup> Calculated with the B3LYP functional. <sup>b</sup> Mean absolute error of the bonds between all atoms apart from C-H bonds. <sup>c</sup> Experimental single-crystal X-ray data from Cambridge Crystallography Data Centre deposition number 798048.

**TABLE S2.** Comparison of error parameters for the DFT optimised geometries of Zr(oxinate)4 in aqueous solution for different functionals with Sapporo-TZP-2012/6-31G(d,p).

| Basis Set<sup>a</sup> | Relative optimisation time | MAE (all)<sup>b</sup>/Å | MAE (Zr-Donor)/Å | MAE (Zr-N)/Å | MAE (Zr-O)/Å | RMSE/Å | CShM |
| --- | --- | --- | --- | --- | --- | --- | --- |
| B3LYP | 1.0 | 0.112 | 0.182 | 0.242 | 0.086 | 0.465 | 1.607 |
| M06-2X | 1.3 | 0.093 | 0.101 | 0.121 | 0.076 | 0.671 | 1.356 |
| ωB97xd | 1.4 | 0.093 | 0.114 | 0.141 | 0.077 | 0.610 | 1.357 |
| PW6B95D3 | 1.1 | 0.091 | 0.107 | 0.134 | 0.070 | 0.839 | 1.778 |
| Experimental XRD<sup>c</sup> |  |  |  |  |  |  | 1.337 |

<sup>a</sup> Calculated with the Sapporo-TZP-2012/6-31G(d,p) basis set. <sup>b</sup> Mean absolute error of the bonds between all atoms apart from C-H bonds.
<sup>c</sup> Experimental single-crystal X-ray data from Cambridge Crystallography Data Centre deposition number 798048.

**Table S3.**
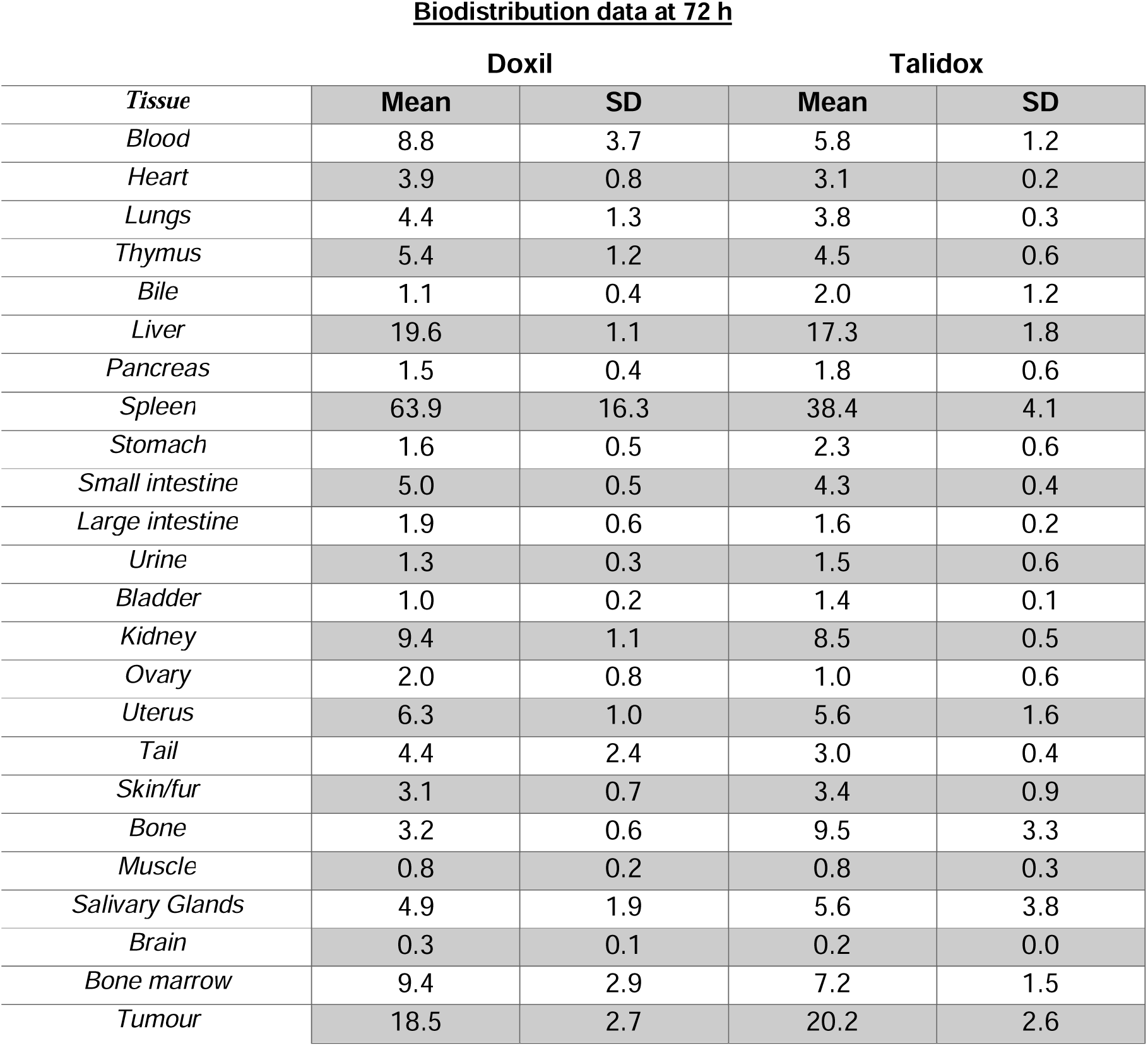
Table compiling the biodistribution data for both formulations, the values represent means and standard deviation of the %IA/g of the organs of interest during the biodistribution at 72 h.

| *Tissue* | **Doxil** |  | **Talidox** |  |
| --- | --- | --- | --- | --- |
|  | **Mean** | **SD** | **Mean** | **SD** |
| *Blood* | 8.8 | 3.7 | 5.8 | 1.2 |
| *Heart* | 3.9 | 0.8 | 3.1 | 0.2 |
| *Lungs* | 4.4 | 1.3 | 3.8 | 0.3 |
| *Thymus* | 5.4 | 1.2 | 4.5 | 0.6 |
| *Bile* | 1.1 | 0.4 | 2.0 | 1.2 |
| *Liver* | 19.6 | 1.1 | 17.3 | 1.8 |
| *Pancreas* | 1.5 | 0.4 | 1.8 | 0.6 |
| *Spleen* | 63.9 | 16.3 | 38.4 | 4.1 |
| *Stomach* | 1.6 | 0.5 | 2.3 | 0.6 |
| *Small intestine* | 5.0 | 0.5 | 4.3 | 0.4 |
| *Large intestine* | 1.9 | 0.6 | 1.6 | 0.2 |
| *Urine* | 1.3 | 0.3 | 1.5 | 0.6 |
| *Bladder* | 1.0 | 0.2 | 1.4 | 0.1 |
| *Kidney* | 9.4 | 1.1 | 8.5 | 0.5 |
| *Ovary* | 2.0 | 0.8 | 1.0 | 0.6 |
| *Uterus* | 6.3 | 1.0 | 5.6 | 1.6 |
| *Tail* | 4.4 | 2.4 | 3.0 | 0.4 |
| *Skin/fur* | 3.1 | 0.7 | 3.4 | 0.9 |
| *Bone* | 3.2 | 0.6 | 9.5 | 3.3 |
| *Muscle* | 0.8 | 0.2 | 0.8 | 0.3 |
| *Salivary Glands* | 4.9 | 1.9 | 5.6 | 3.8 |
| *Brain* | 0.3 | 0.1 | 0.2 | 0.0 |
| *Bone marrow* | 9.4 | 2.9 | 7.2 | 1.5 |
| *Tumour* | 18.5 | 2.7 | 20.2 | 2.6 |

